# Functional Ultrasound Imaging Reveals Activation Properties of Clinical Spinal Cord Stimulation Therapy Stimulation Programming

**DOI:** 10.1101/2024.10.22.619739

**Authors:** Koeun Lim, P Sean Slee, Andrew Kibler, Steven Falowski, Kasra Amirdelfan

**Affiliations:** BIOTRONIK NRO Inc., Lake Oswego, OR, USA; Neurosurgical Associates of Lancaster, Lancaster, PA, USA; Boomerang Healthcare, Walnut Creek, CA, USA

**Keywords:** Spinal cord stimulation, functional ultrasound imaging, spinal blood volume, hemodynamic response

## Abstract

**Objectives:** Spinal cord stimulation (SCS) therapy has long been established as an effective treatment for chronic neuropathic pain. However, methodological limitations have prohibited the detailed investigation of the activation patterns produced in the spinal cord during therapy. Functional ultrasound (fUS) is an emerging technology that monitors local hemodynamic changes in the brain that are tightly coupled to neural functional activity [1–3]. Previous studies have demonstrated that the high sensitivity and spatiotemporal resolution of fUS can be used to monitor activation in the spinal cord [4, 5]. In this study, fUS was used to investigate neuromodulation patterns produced by clinical SCS paradigms in an ovine model that enabled testing with implanted clinical hardware.

**Materials and Methods:** Activation of local dorsal horn regions during SCS therapy was evaluated using fUS to detect hemodynamic changes in flowing spinal blood volume (ΔSBV). Briefly, male ovine subjects were anesthetized and laminectomies were performed at T12-L1 to expose the spinal cord. The spine was mechanically fixed to reduce breathing-induced motion. Standard SCS leads were percutaneously implanted midline overlying the dura of the exposed cord to enable stimulation and recording. A second lead was implanted in the epidural space anterior to the laminectomy for recording evoked compound action potentials (eCAPs). Motor thresholds were determined from electromyographic (EMG) signals recorded from subdermal needle electrodes. Hemodynamic activation patterns produced by SCS therapy were mapped across two vertebral segments in the superficial dorsal horn (SDH) at amplitudes between 100%-200% eCAP threshold (eCAPT). The magnitude and volume of significant ΔSBV during SCS was quantified and compared across conditions.

**Results:** eCAP and motor thresholds varied widely between different representative SCS programs in current clinical use. Compared with motor threshold, we found eCAPT to be a more stable and reliable ratiometric reference to establish neural drive from stimulation, consistent with previous literature [6]. SCS stimulation resulted in significant activation of the SDH in differing patterns across two vertebral segments. The magnitude and volume of ΔSBV increased at higher amplitudes and was typically maximal in the SDH regions underlying the active electrodes. Activation persisted for several seconds following SCS therapy cessation, suggesting the engagement of neurophysiological processes with correspondingly long time constants. In addition, therapy mode significantly influenced total area and depth of ΔSBV. Multiphase therapy produced a larger area of ΔSBV that extended deeper into the spinal cord relative to single phase therapies.

**Conclusions:** This work demonstrates that fUS can effectively measure SCS neural response patterns in the pain processing laminae of a large animal model implanted with a clinical SCS system. Hemodynamic responses in the spinal cord varied significantly across SCS therapy modes, with Multiphase stimulation providing a greater area of coverage and depth of response versus other common stimulation types.

## INTRODUCTION

Spinal cord stimulation (SCS), a neuromodulation therapy used in the management of chronic neuropathic pain, emerged in 1967, shortly after the seminal paper on the gate control theory of pain perception [7]. Traditionally, SCS is employed using biphasic, rectangular pulse trains which contain one therapeutic phase (single phase) coupled with a non-therapeutic charge balancing phase, each repeating at frequencies ranging from 20 Hz–1.2 kHz [8–10] and in some cases up to 10 kHz. This mode of stimulation is delivered at amplitudes believed to activate large sensory Ab fibers in the dorsal column (DC), which in turn antidromically activate inhibitory networks in the dorsal horn and suppress pain transmission [11–13]. Activation of DC fibers also produces a tingling sensation, commonly referred to as paresthesia. A sub-perception (paresthesia-free) mode of SCS has more recently been demonstrated to achieve high efficacy across a broad range of parameters [9, 14–18]. This mode of stimulation is delivered below the action potential threshold of DC fibers, hypothesized to modulate excitability and synaptic coupling locally in the dorsal horn [19–23] and typically acquires maximal efficacy over hours-days consistent with neuroplasticity timescales [24, 25].

Spinal cord stimulation (SCS) therapy has long been established as an effective treatment for chronic neuropathic pain. Differences between the mechanisms of action engaged by clinically available SCS therapy modes have been claimed based on studies utilizing brain imaging [26, 27], electrophysiology [28], genetics [29, 30], proteomics [31] and computational modeling [32]. However, methodological limitations have prohibited a detailed investigation of the activation patterns produced in the spinal cord grey matter during therapy.

Functional ultrasound (fUS) is a newly established technology that monitors local hemodynamic changes in the brain and spinal cord that are tightly coupled to the underlying neural activity [1–3, 5, 33]. Specifically, local spinal blood volume has been shown to increase monotonically with increases in the firing rate of neuronal populations within that region [3]. Previous studies have demonstrated that the high sensitivity and spatiotemporal resolution of fUS can be used to detect regions of altered neural activity in the spinal cord during activation of nociceptive input [5] and SCS [4]. In this study, fUS was used to illuminate dorsal horn neural activity responses to neuromodulation patterns produced by single phase and multi-phase clinical SCS paradigms in an ovine model with clinical stimulation hardware.

## MATERIALS AND METHODS

### 2.1 Animal preparation procedure

Experimental procedures were approved by the Legacy Research Institute Animal Care and Use Committee. The animals we cared for according to the USDA Animal Welfare Act standards and the eighth edition of *The Guide for Care and Use of Laboratory Animals*. Male ovine subjects (N=7) of the Polypay breed we used in this study. The animals were 18-36 months of age and weighed between 60-75 kg. Prior to the experimental procedure the animals were housed in a colony in a temperature and humidity-controlled environment with a 12 hour light/dark cycle with food provided 2 times/day and water provided ad libitum.

On the morning of the experiment, the subject was premedicated with midazolam (0.35 mg/kg) and meloxicam (0.2 mg/kg) prior to transportation to the procedure room. An intravenous (IV) catheter was placed in the right saphenous vein followed by induction of anesthesia with ketamine (20 mg/kg IV) and midazolam (0.25 mg/kg IV, given in 25% increments to effect). Next the subject was intubated with a #11 endotracheal tube and placed in the prone position on the procedure table. Respiration with a mechanical ventilator (Hallowell 2002IE) was used to maintain tidal volume (7-10 ml/kg) using a 95% O_2_/CO_2_ mixture combined with Isoflurane (1-4%) to effect.

General anesthesia was maintained with ketamine (0.4-4 mg/kg/hr by constant rate infusion, CRI), midazolam (0.1-0.9 mg/kg/hr CRI), fentanyl (0.0001-0.005 mg/kg/hr CRI) and isoflurane (0.5-1%). Hetastarch (1 ml/kg/hr IV) was given to maintain circulating blood volume and saline given for hydration (1 ml/kg/hr IV). Core body temperature was maintained using a heating blanket that circulated warm air (Bair Hugger Model 750; 3M). Vitals including heart rate, SpO2, CO_2_, respiration rate (15-22 bpm), blood pressure and core temperature were monitored throughout the procedure. Depth of anesthetic plane was assessed by jaw tone every 15 minutes.

Once the subject was in a deep anesthetic plane, a tuohy needle was placed in the epidural space between T5-T6 under fluoroscopic guidance to allow placement of an epidural catheter for local anesthesia (bolus delivery of 10% lidocaine, 0.5 ml/30 min). A set of control experiments were conducted to confirm epidural block at T5-T6 did not influence SCS or eCAP recording at T12-L1 (see Section 2.3 below for details).

Next a laminectomy was performed between the 13^th^ thoracic vertebra and the first lumbar vertebra (T13-L1, in most animals) to expose the spinal cord for fUS imaging. Compared with more superior spinal levels, this ovine spinal level was chosen because of its shorter spinal processes, which facilitated positioning of the ultrasound transducer. To reduce breathing motion the spine was mechanically fixed by wire to a Tufail Bookwalter Retractor System (BR18-77000) coupled to the procedure table. This configuration resulted in total displacement of the fUS image of 0.5-1 mm during the respiration cycle. A second tuohy needle was placed at L2/L3 junction to implant a percutaneous octal SCS lead (BIOTRONIK Resilience, 3 mm electrode/4 mm spacing) midline overlying the dura of the exposed cord to enable stimulation. Consistent with recent modeling studies, eCAP thresholds were initially elevated for stimulation over the laminotomy region relative to other regions due to dorsal leakage of stimulation current that is normally insulated by ligament and bony spinous structures [34]. To enable more direct comparison with standard clinical configurations the dorsal half of each cylindrical electrode was insulated with a thin coating of acrylic. A custom paddle lead (8 electrodes, 3x1 mm, 5 mm spacing) was implanted midline in the epidural space superior to the laminectomy (T12) for recording evoked compound action potentials (eCAPs) from the dorsal column. Pairs of subdermal needle electrodes were placed at 16 medial and lateral positions on both sides of the animal from the shoulder to hind quarter to enable recording of electromyographic (EMG) signals. Finally, a subcutaneous ground reference wire was placed on the right side of the subject.

### 2.2 Clinical Spinal Cord Stimulation Hardware

Stimulation was delivered to the spinal cord using a clinical-grade external pulse generator (EPG) [35]. The EPG is battery powered, has 16 channels, and is programmed and controlled via Bluetooth by a computer running validated software. The EPG was connected to the implanted SCS lead using a custom adaptor.

### 2.3 Electrophysiological Recordings

eCAPs were recorded from the dorsal column at multiple locations rostral to the stimulation and imaging site using the custom paddle lead (see **Figure 1**). Each recording channel consisted of one electrode referenced to one or more electrodes on the same paddle lead. Signals were filtered (0.1-3 kHz) and differentially amplified (gain = 2k; Bio-Amp BMA-400) prior to recording using a A/D board (20 kHz; NI USB X Series USB) running custom software. An input channel was used to record the stimulation artifact to synchronize the eCAPs with the stimulation pulses. EMG was recorded from the subdermal needles using a commercial intraoperative neuromonitoring system (Nicolet, Endeavor).

**Figure 1.**
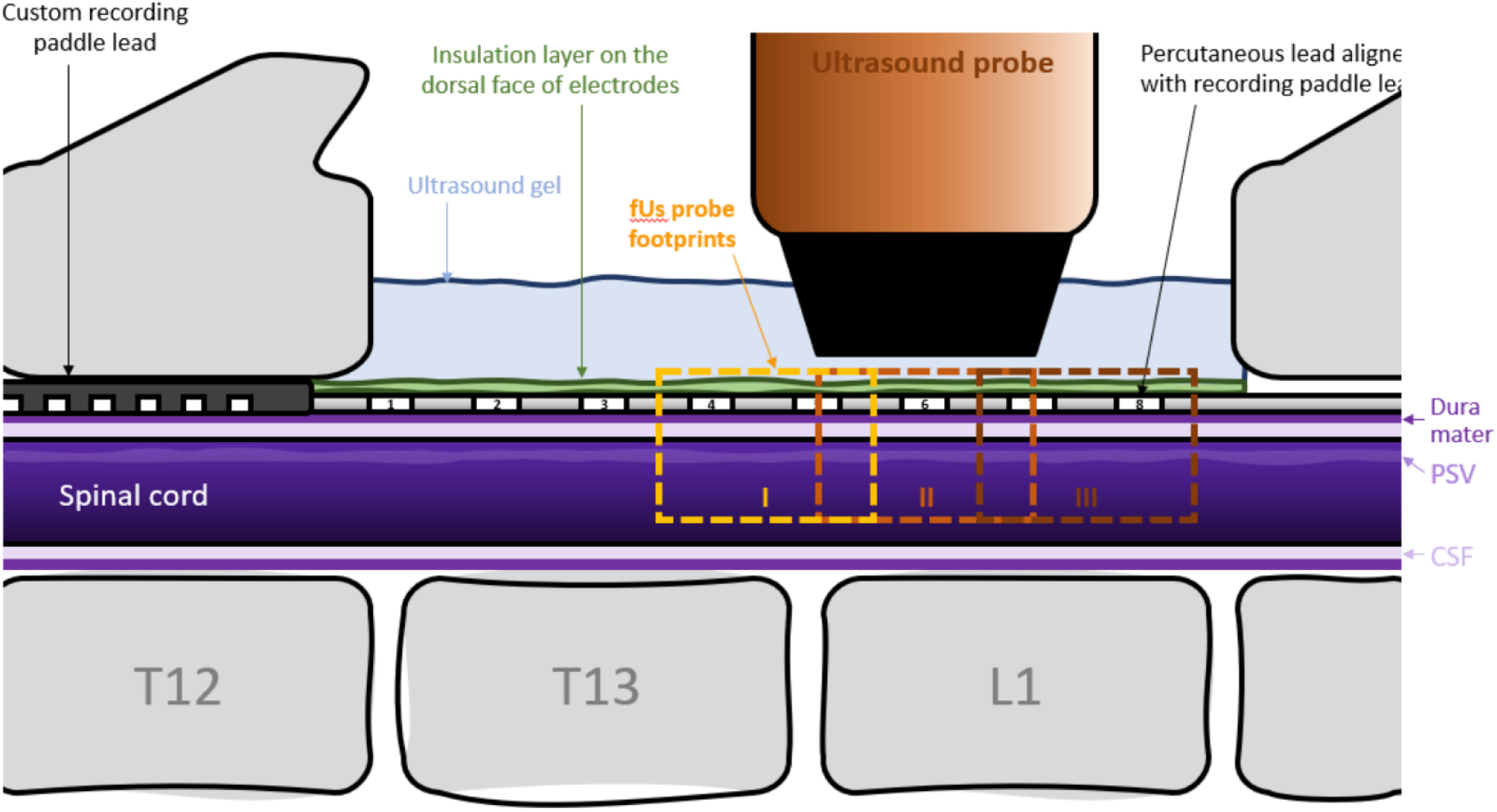
Experimental Configuration. A standard octal lead placed midline covering laminectomy and a custom paddle for recording ECAPs. The fUS probe was placed dorsal to the left SDH to monitor hemodynamic responses to SCS in a region proximal to the central active contact (orange box II) or in one of the adjacent distal regions (boxes I and III). Posterior spinal vein (PSV), cerebrospinal fluid (CSF)

### 2.4 Functional Ultrasound Imaging

The dural surface was prepared for imaging as described previously [4]. The ultrasonic probe, which was fixed on a 3-axis motor system, was then positioned in the coronal plane just above the center of electrode 6 on the SCS lead (see **Figure 1**) using the Icoscan live acquisition software (Iconeus, Paris, France). After alignment the probe was moved lateral to center it above the left dorsal horn (Y=∼-2 mm) which was identified using the posterior spinal vein (PSV) as a landmark [5].

Functional ultrasound imaging was performed using a linear ultrasound probe (128 elements, 15 MHz, 110 mm pitch, 8 mm elevation focus, Vermon) driven by an ultrafast ultrasound scanner (Verasonics, Kirkland, WA: 128 channels, 62.5 MHz sampling rate), which was driven with Icoscan live. The fUS imaging sequence operated as follows: the spinal cord was insonified by 10 successive tilted plane waves with angles varying from 210° to 10° with a 5.5 kHz pulse repetition frequency. The backscattered echoes were recorded by the transducer array and beamformed to produce a block of 200 consecutive ultrafast images with a framerate of 500 Hz. To filter the spinal blood volume (SBV) in the 200 frame block, the tissue signal was removed by applying a clutter filter based on singular value decomposition (SVD) [36]; the 60 first singular vectors were removed, as they correspond mainly to the tissue space. Finally, a Power Doppler image was obtained by integrating the energy of the filtered frames, resulting in a Power Doppler image every 400 ms (2.5 Hz frame rate).

### 2.5 Clinical Spinal Cord Stimulation Therapy

Five different clinical SCS therapies were programmed to the EPG using clinical software [35]; Burst (BST), Multi-frequency (MF), High kHz frequency (HF), Multiphase (MP) and Traditional (Trad) as detailed in Table 1. The amplitude resolution of the EPG was 0.1 mA. If finer resolution was needed for a given animal/therapy, all active electrodes were connected to each other via a resistor network (100, 121 or 200-ohm) using custom hardware prior to adapting to the SCS lead.

**Table 1.** Clinical SCS therapy parameters evaluated in this study. Active anodes and cathodes numbers correspond to contacts on a typical octal lead with contact 1 being distal. The right side of the table lists the numbers of subjects or locations for which fUS, eCAPT and MT were evaluated for each therapy type.

| Therapy Mode | Frequency (Hz) | Pulse Width (μs) | Active Cathodes | Active Anodes |  |  |
| --- | --- | --- | --- | --- | --- | --- |
| Control (No SCS) | - | - | - | - |  |  |
| Burst | 500 (40) | 1000 | 6 | 7 |  |  |
| Multi-Freq | 50 / 1200 | 300 / 150 | 3 / 6 | 2 / 7 |  |  |
| High kHz Freq | 8333 | 30 | 6 | 7 |  |  |
| Multiphase | 300 | 300 | 4, 6, 8 | - |  |  |
| Traditional | 50 | 300 | 6 | 5, 7 |  |  |
| Therapy Mode | N Subs fUS | N Subs eCAPT | NSubs MT | fUS N Prox Locs | fUS N Dist Locs | fUS N Total Locs |
| Control (No SCS) | 7 | - | - | 37 | 29 | 66 |
| Burst | 7 | 7 | 2 | 18 | 19 | 37 |
| Multi-Freq | 7 | 7 | 3 | 19 | 19 | 38 |
| High kHz Freq | 5 | 6 | 3 | 14 | 23 | 37 |
| Multiphase | 7 | 7 | 4 | 40 | 26 | 66 |
| Traditional | 4 | 5 | 1 | 13 | 13 | 29 |

### 2.6 Experimental Procedure

In each experiment the imaging location (proximal or distal to the common cathode) and SCS therapy type were block-randomized. Specifically, all therapies were tested in random order at one location prior to moving the probe to a new location. Prior to SCS a series of control images were collected in the first location. Next the eCAP threshold (eCAPT) was determined for each therapy. Stimulation was initiated at 0.1 mA and incremented by 0.1 mA every 2-3 seconds using 2 second onset/offset ramps between level transitions. Stimulation was stopped when either a clear eCAP or EMG response was observed in steady-state period following amplitude transitions. eCAPT was initially identified visually using an oscilloscope then confirmed post-hoc using custom analysis software written in MATLAB.

Each therapy was initially tested at an amplitude equivalent to 100% eCAPT. For each fUS scan 40 seconds of baseline preceded 20 of SCS. The pattern was repeated twice with additional 40 seconds of off time at the end for a total of 220 seconds of data collection (see **Figure 2**). The therapy amplitude was incremented by 10% and another scan collected. This was repeated until reaching motor threshold or 200% eCAPT. Control scans (no SCS) of the same duration were collected between testing different therapies. Once all therapies in a location were tested, the probe was moved to a new location. Therapies were again tested in a new randomized order. In most experiments, 3-4 hours of fUS imaging could be completed prior to decline in signal quality indicated by a decrease in the intensity and contrast of the control scans.

**Figure 2.**
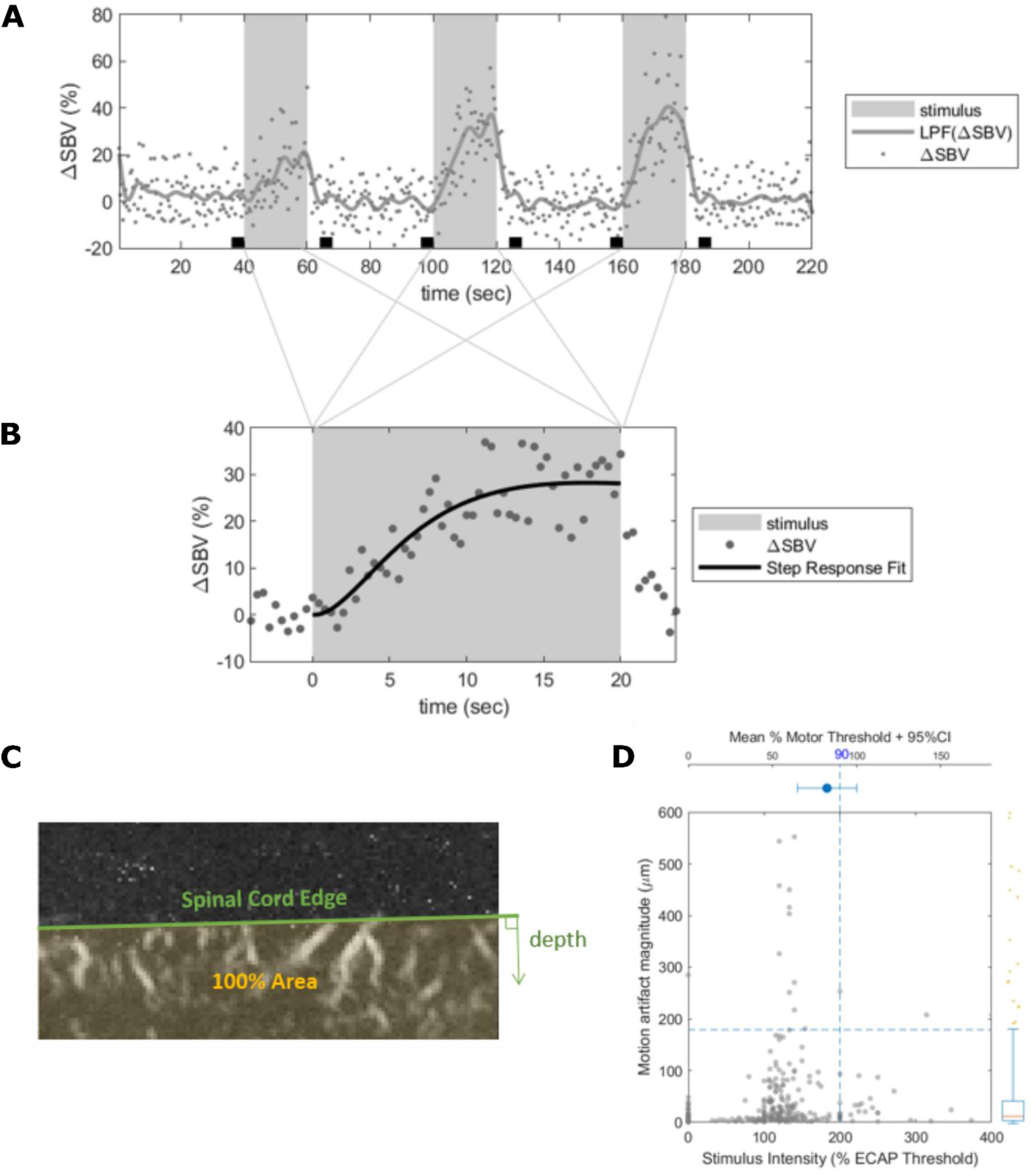
A, Example of hemodynamic responses induced by 3 bouts of multiphase SCS therapy indicated by gray bars. Individual points indicate the instantaneous change in spinal blood volume (DSVB) measured in a single region of interest. The solid gray line shows the lowpass filtered response. The black boxes indicate the pre-stimulus intervals used to measure the baseline SBV. B, The points show the response in A averaged over the three SCS presentations. The solid black line shows the step response curve fit used to extract the steady-state response. C, The green line indicates the boundary of the dorsal edge of the spinal cord used to measure depth of activation. D, Scatterplot of motion artifact magnitude versus stimulus amplitude used to define data inclusion criteria. Data in the lower left box were included for further analysis.

### 2.7 fUS image processing

fUS image sequences contain both spatial and temporal information about neural activation. Hence, proper treatment in both spatial and temporal domains is needed to accurately infer the location and magnitude of neural activation induced by neuromodulation. Each pixel in fUS scans contain time-varying power doppler measurements that reflect the change in spinal blood volume (ΔSBV). To characterize ΔSBV at each pixel at a fixed reference frame, motion-correction was applied using intensity-based image registration to adjust for procedural motion artifacts such as breathing, drifting, and muscle movements (Image Processing toolbox in MATLAB). Motion artifact estimates were obtained during the registration process to monitor the quality of the data.

Once the registration was completed, the surface edge of the spinal cord was semi-automatically defined in the fixed reference frame. These spinal cord edges served two purposes: (1) to constrain the signal processing and analysis within the spinal cord, eliminating noise from the surrounding external tissues and debris in the gel; and (2) to serve as the reference point for the depth definition.

Within the spinal cord pixels, temporal signal processing was performed for each pixel. Because of the long in-vivo imaging duration (150∼220 seconds), fUS time-series are prone to slow drift and offset caused by Brownian noise and hysteresis. Temporal data processing removed such slow drifting by correcting for pre-stimulus and post-stimulus offsets. Pre-stimulus interval was defined as -20% of the stimulus bout duration relative to the stimulus onset (t_on = 0 sec). Post-stimulus interval was defined as +20% to +40% of the stimulus bout duration relative to the stimulus offset (t_off = dur_stim). The same pre-stimulus interval was used to determine the baseline spinal blood volume. In accordance with earlier fUS studies [3, 5], ΔSBV at time t relative to t_on was defined relative to this baseline spinal blood volume such that 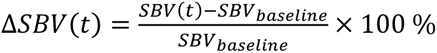. Since each scan contains multiple SCS bouts, these repeated Δ*SBV*(*t*) measurements are averaged across the bouts so that there is single Δ*SBV*(*t*) time-series per pixel per scan. Then the time average during SCS was taken per pixel 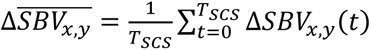 as the single-point descriptive summary statistics of the hemodynamic response to obtain the spatial response map, where (*x*, *y*) indicates the pixel coordinate. Resulting 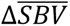 maps are then aggregated per animal across all pixels in all scans to find an activation threshold, which can be described as the pixels with 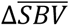 significantly greater or less than 0 at *α* = 0.05. Here, the threshold is found for each animal to account for the individual sensitivity to SCS and anesthesia, but across therapy modes including control to compare therapy effects.

### 2.8 Analysis of DSBV response area, depth and magnitude

The collection of pixels with significant mean response is called region of interest (ROI). Depending on the polarity of 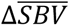, two types of ROI can be defined. In vascular physiology, positive blood volume change is called hyperemia, and negative blood volume change is called congestion. Hence, the collection of pixels with positive 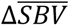 indicates hyperemia, termed ROIh, while the collection of pixels with negative 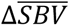 indicates congestion, termed ROIc. For each of ROIh and ROIc, hemodynamic response was quantified in both spatial and temporal domains. In spatial domain response area and depth were quantified, and in temporal domain Δ*SBV* magnitude was quantified.

To quantify Δ*SBV* magnitude, the collection of Δ*SBV*(*t*) within the identified ROIh and ROIc were first averaged to obtain mean ROI hemodynamic response time-series 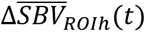 and 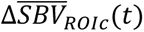. Then the hemodynamic response parameters were estimated by fitting a second-order low-pass step response function such that 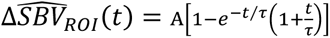. Here, A is the response amplitude where *A* > 0 would be associated with ROIh and *A* < 0 with ROIc.

Differences in the tissue volume captured in fUS images across scans were accounted for by taking the percentage of the number of pixels within each ROI (*n*_*ROI*_) out of all spinal cord pixels (*n*_*SC*_) such that 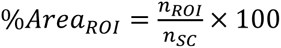. For depth, 75^th^ percentile of the distance between the surface of the spinal cord and ROI pixels was calculated (**Fig 2C**).

When quantifying hyperemic ROI, because of the statistical threshold applied to fUS images to classify responsive vs. non-responsive pixels, the response features inherit the same bimodality. In other words, significant hemodynamic responses are not generally observed on every trial at a fixed stimulus level. This trait manifests as bimodal distribution of ROI features, one peak at zero and another peak at a non-zero value. Such bimodality is prone to biased estimate of the mean response especially for small sample size where uniform sampling is procedurally limited by the animal’s physiological sensitivity to anesthesia and experimental procedures. To correct for the sampling bias, response probability as a function of stimulus intensity was obtained for each of the therapy mode by fitting a Gaussian cumulative distribution function to the binarized ROI responses. First, for each therapy per subjects, ROI responses are grouped according to stimulus intensity bin of 20% eCAPT. The proportion of *n*_*ROI*_ > 0 is obtained, which is then pooled across subjects to fit the activation function that maps expected response probability as a function of stimulus amplitude such that 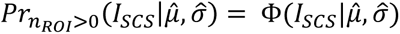 where Φ is a cumulative Gaussian function with estimated mean *μ̑* and variance *σ̑*^2^. The hat symbol ^ indicates maximum likelihood estimate. This probability function translates bimodal ROI response (zero or non-zero) to continuous numerical space by scaling non-zero responses to predict expected ROI responses, which can be expressed mathematically as 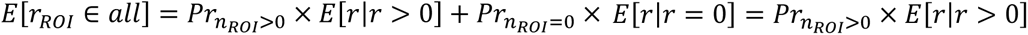 Here, *E*[*r*_*ROI*_ ∈ *all*] is the expected ROI response considering all samples including both zero and non-zero responses, *E*[*r*|*r* > 0] is the expected non-zero ROI responses, and *E*[*r*|*r* = 0] is the expected zero responses that reduce to 0. For instance, for high stimulus intensity, the response probability approaches 1 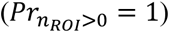, meaning the average non-zero response magnitude would equal the expected response magnitude. On the other hand, when given low stimulus intensity where the response probability is only 0.1 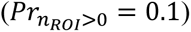, although the average non-zero response magnitude may be X (*E*[*r*|*r* > 0] = *X*), it is only observed 1 out of 10 times; therefore, the expected response for both zero and non-zero response would be 0.1X. In summary, response probability functions act as a weighting function to scale non-zero ROI responses in order to predict expected ROI responses encompassing both hemodynamic response modes (zero vs non-zero).

In 7 animals, 323 scans were obtained with corresponding eCAPT measurements. Some motor thresholds were not measurable, in which case mean motor-to-eCAP threshold ratio was used to limit SCS amplitude. First, the upper 95% confidence interval of the mean motor-to-eCAP threshold ratio was found, which corresponded to 222% eCAPT stimulus amplitude (**Fig. 2D**, top subplot), then 90% of this upper bound was estimated to be at 200% eCAPT. In other words, SCS amplitude was limited to 90% of the motor threshold average’s upper bound (**Fig. 2D**, bottom left box). In addition, the scans with motion artifact exceeding 95^th^ percentile were rejected. The median motion artifact was 0.13 pixels (equivalent to 13μm) and the 5% outliers greater than 1.79 pixels (equivalent to 179μm) were rejected. The resulting total data rejection rate was 11.6%. **Figure 2D** illustrates the data inclusion criteria (bottom left box).

### 2.8 Statistics

Repeated-measures marginal ANOVA was performed on the quantified hemodynamic response features to observe effects of experimental conditions such as therapy mode, stimulus intensity, and probe location relative to the main cathode, as well as procedural artifacts including order effect and motion artifact. Once the significant marginal factors were identified, linear mixed-effects models (LMEs) were fit to investigate the relationship between the experimental conditions and the hemodynamic responses, weighted by the activation function. In short, with control data serving as the anchor, the slopes of the linear relationship between the response and the stimulus amplitude for all therapy modes were calculated with subject and motion artifact as the random effect. Pairwise comparisons were then performed for the gain (slope) parameters between each pair of therapy modes. Bonferroni-Holms correction was applied to adjust for the multiple comparisons. All statistical analyses were performed in MATLAB (statistics toolbox). The statistical methods and results presented in this manuscript were performed based on the consultation with a statistician in the BIOTRONIK clinical department.

## RESULTS

Activation of the spinal cord during SCS therapy was evaluated using fUS imaging to detect hemodynamic changes in spinal blood volume in the region of the dorsal horn. Five different clinical SCS therapies were tested in an ovine model implanted with clinical SCS hardware (see **Table 1**). Hemodynamic responses to SCS therapy were mapped in a parasaggital plane containing the dorsal horn in 3 rostro-caudal regions spanning two vertebral segments. SCS amplitudes between 100%-200% eCAP threshold were included in the analysis (see **Figure 1**).

### 3.1 fUS demonstrates hemodynamic responses to clinical SCS below motor threshold

We measured hemodynamic changes in spinal blood volume for each SCS therapy type and level. For each scan three bouts of SCS were presented for 20 s with 40 s of rest between each presentation (**Fig. 2A**). In most cases significant DSBV below eCAPT were not seen. As stimulation amplitude increased between 100-150% eCAPT, significant DSBV were detected. A typical single-voxel hemodynamic response is shown in **Figure 2B** for multiphase SCS therapy at a level of 111% eCAPT (**red box voxel in Fig. 3B**). Responses tended to build up over ∼10 s relative to stimulus onset before reaching a plateau, corresponding to ∼20-30% increase in SBV in the example shown (**Fig. 2B**). Response patterns across stimulus bouts were typically consistent with some showing a tendency for a greater response to the second and third SCS presentation.

**Figure 3.**
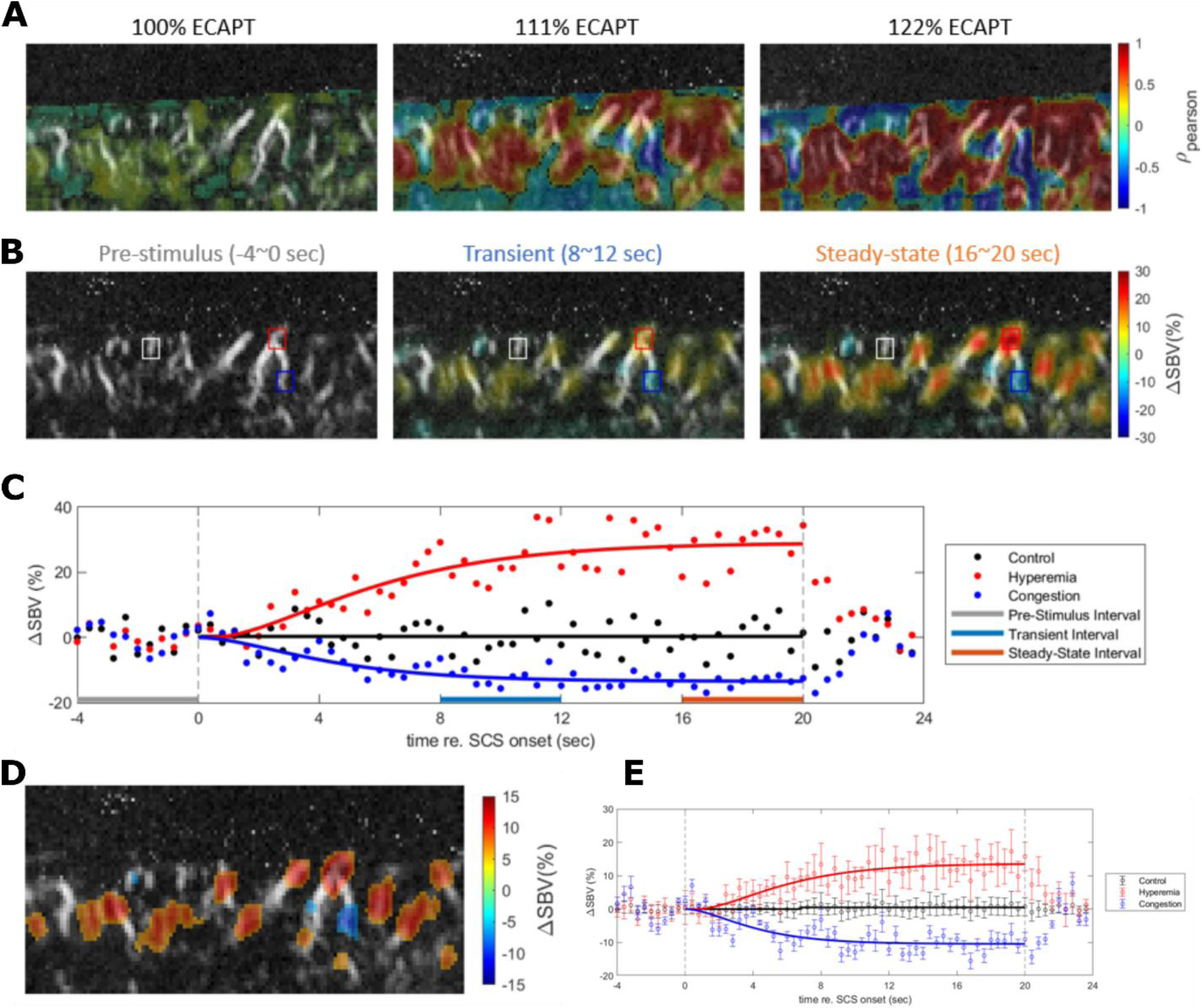
Example of hemodynamic responses to SCS of increasing amplitude. A, Cross correlation of multiphase SCS on-time versus DSBV for increasing SCS amplitude (left to right panels). B, DSBV in response to multiphase SCS for increasing SCS amplitude (left to right panels). C, Time-course of DSBV measured in the ROIs indicated by the colored squares in B for the middle panel at 111% eCAPT. Points indicate raw data and solid lines are step-response curve fits as described in Figure 2. Gray, blue and red thick bars indicate the pre-stimulus baseline, transient and steady-state intervals, respectively. D, DSBV in response to multiphase SCS at an amplitude of 111% eCAPT after thresholding for only statistically significant DSBV. E, Mean and SD of DSBV across all positive (red, hyperemia) and negative (blue, congestion) ROIs shown in D. Black (control) show responses from all non-significant pixels.

Before quantifying bout-by-bout responses, motion artifacts (**Fig. 2D** vertical axis) and the correlation between the stimulus and power doppler signal (correlation maps in **Fig. 3A**) were first estimated on a per-scan basis using the entire time-series per scan. To avoid rapid transients near onset and offset, stimulus-off interval defined at 60% of the stimulus OFF duration before each stimulus onset was used, and stimulus-on interval defined at 60% of the stimulus ON duration before each stimulus offset to calculate the correlation between stimuli and responses. In an example in Figure 3C, stimulus-off interval corresponds to -12 ∼ 0 seconds, and stimulus-on interval corresponds to 8∼20 seconds in x-axis.

Quantification of the hemodynamic response’s temporal dynamics required a data processing approach that accounted for slow drifts (see Materials and Methods) that impact the magnitude of the hemodynamic response. Baseline correction was performed for individual bouts, and then the average response was obtained across all bouts within the scan (**Fig. 2B**). Then a second order low-pass step response function was fitted to the average response (solid black trace in **Fig. 2B**, see Materials and Methods). The magnitude of the response was taken as the plateau amplitude of the fit which corresponded to a DSBV of 28% for the example shown. As stimulation amplitude increased to motor threshold, fUS image displacements, which correlated with stimulus onset, were noted. Small shifts within the image plane (<1.79 pixels equivalent to 179μm) were corrected while scans with larger shifts were rejected from the analysis (**Fig. 2D** vertical axis, see Methods).

### 3.2 Hemodynamic responses reveal hyperemia and congestion independent of motor activation and respiratory motion

Examples of the spatio-temporal properties of the hemodynamic responses to clinical SCS are shown for multiphase therapy in **Figure 3**. Correlation maps showing the cross-correlation between stimulus and response for the three stimulus levels (100%, 111%, and 122% eCAPT) are shown in **Figure 3A**. At 100% eCAPT (left) both positive correlations indicating regions of hyperemia (DSBV increases), and negative correlations indicating regions of congestion (DSBV decreases) are readily apparent in **Figure 3A**. As stimulus intensity increases (middle, right) the strength of the correlations increases while the spatial pattern and polarity remains largely unchanged. In order to observe temporal dynamics, snap shots of DSBV contour maps were taken in response to 111% eCAPT stimulus at different times relative to the SCS onset (Figure 3B). **Figure 3B** shows contour maps corresponding to pre-stimulus, transient, and steady-state intervals (marked with horizontal bars in **Figure 3C**). Similar to the correlation maps at different intensities (**Fig. 3A**), the spatial distribution of hyperemia and congestion is preserved during the stimulation while the magnitude of the response increases towards steady-state. Temporal dynamics (**Fig. 3C**) was further examined in the voxels corresponding to the maximum hyperemia and congestion relative to a control voxel with no hemodynamic response (**Fig. 3B**). As seen in the example shown in **Figure 2**, the DSBV hemodynamic response builds over about the first 12 s and then reaches a steady-state (**Fig. 3C**). This temporal pattern was similar for both increases and decreases in SBV.

In order to aggregate spatial and temporal response parameters per scan, hemodynamic responses were averaged within ROI defined by the activation map (**Fig. 3D**). **Figure 3D** shows an example activation map that includes hyperemia (warm pixels) and congestion (cool pixels) significantly greater than 0 (alpha = 0.5, pooled across all scans per animal). Average hyperemia DSBV (red markers, mean ± SD) and congestion DSBV (blue markers, mean ± SD) time-series are shown in **Figure 3E** against the out-of-ROI DSBV control time-series (black markers, mean± SD).

### 3.3 Hemodynamic responses varied across Clinical SCS Therapy Modes

Hemodynamic responses to different SCS therapies were measured over a range of amplitudes scaled relative to eCAP threshold. **Figure 4A-D** shows examples of responses in one subject (#106-152) at comparable amplitudes with the fUS transducer positioned above the dorsal horn in the proximal imaging region (see Methods). The PSV is apparent near the dorsal edge of the SC at the top of the image (arrow in **Fig. 4A**). Traditional therapy produced no significant activation in the SC (**Fig 4A**). In contrast, both Multiphase (**Fig. 4B**) and Burst (**Fig. 4C**) SCS therapy activated a large area of the SC that extended deep into the SC. High-kHz SCS also produced strong activation although in a more restricted area in the dorsal region of the SC (**Fig. 4D**). **Figure 4E,F** shows examples of responses to Multi-Frequency and Multiphase therapies, respectively, measured from a different subject (#107-69). Here large areas of activation extend deep into the SC for both therapy types and both hyperemia and congestion were detected. However, the magnitude of the response was greater for Multiphase compared to Multi-Frequency.

**Figure 4.**
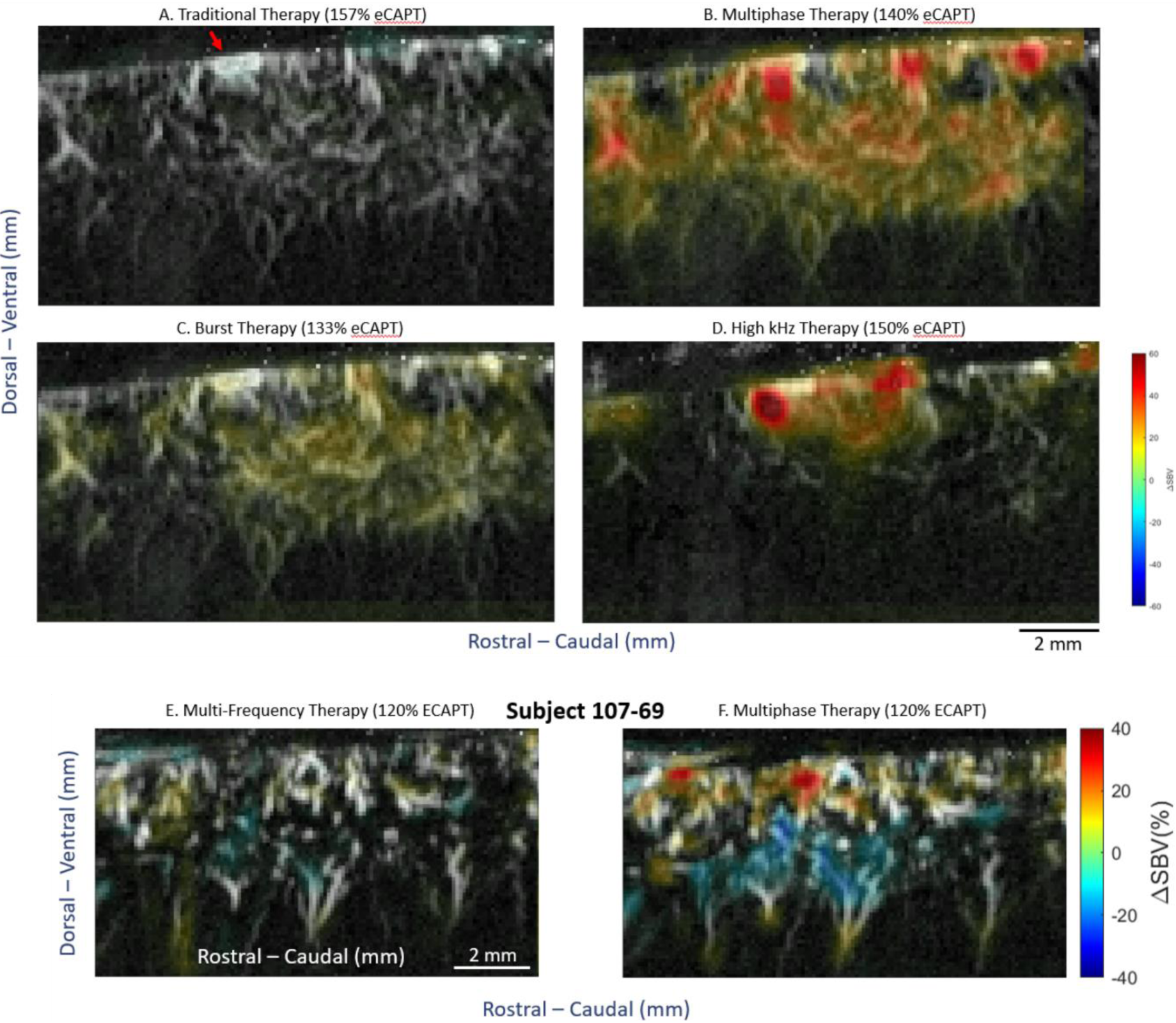
A-D, Examples of SC activation patterns induced by different SCS therapies (sub-106-152). The fUS transducer was positioned in the proximal region near the centermost cathode and 1.6 mm lateral to the midline (at the level of the dorsal horn). The DSVB induced by SCS (color map) is superimposed over the vascularization of the spinal cord (black and white background). A, Traditional. B, Multiphase. C, Burst. D, High kHz. E-F, Examples of SC activation patterns induced by multi-frequency and multiphase SCS recorded from a different subject (107-069).

In order to quantify hemodynamic responses across therapy modes, fUS responses were pooled across subjects, SC locations and amplitudes and fit with a LME model using therapy mode and SCS amplitude as fixed effects (see Methods). **Figure 5A** shows the resulting hemodynamic activation probability for each therapy mode as a function of SCS amplitude (scaled by eCAPT, **Fig. 5B**). The activation function for Traditional therapy (right panel) had a relatively shallow slope with modest peak probability reflecting the fact that traditional therapy only produced significant hemodynamic responses in a few cases. In contrast, the activation functions for Burst, MF, HF and MP had steeper slopes with a rapid rise around 100% eCAPT that peaked at high probabilities of activation. This reflects the fact that these therapies typically drove hemodynamic responses at amplitudes above eCAPT. The activation functions were next used to quantify the differences in the features of the hemodynamic responses across therapy modes. Quantification of three hyperemia features (DSBV increases) were evaluated because previous studies have demonstrated a direct correlation between hyperemia and neural activation through the mechanism of neurovascular coupling [3]. These features were hyperemia Magnitude (peak DSBV), %Area (fraction of SC with significant positive DSBV) and Depth (75^th^ percentile of the distribution of DSBV depths relative to the dorsal surface, see Methods for details).

**Figure 5.**
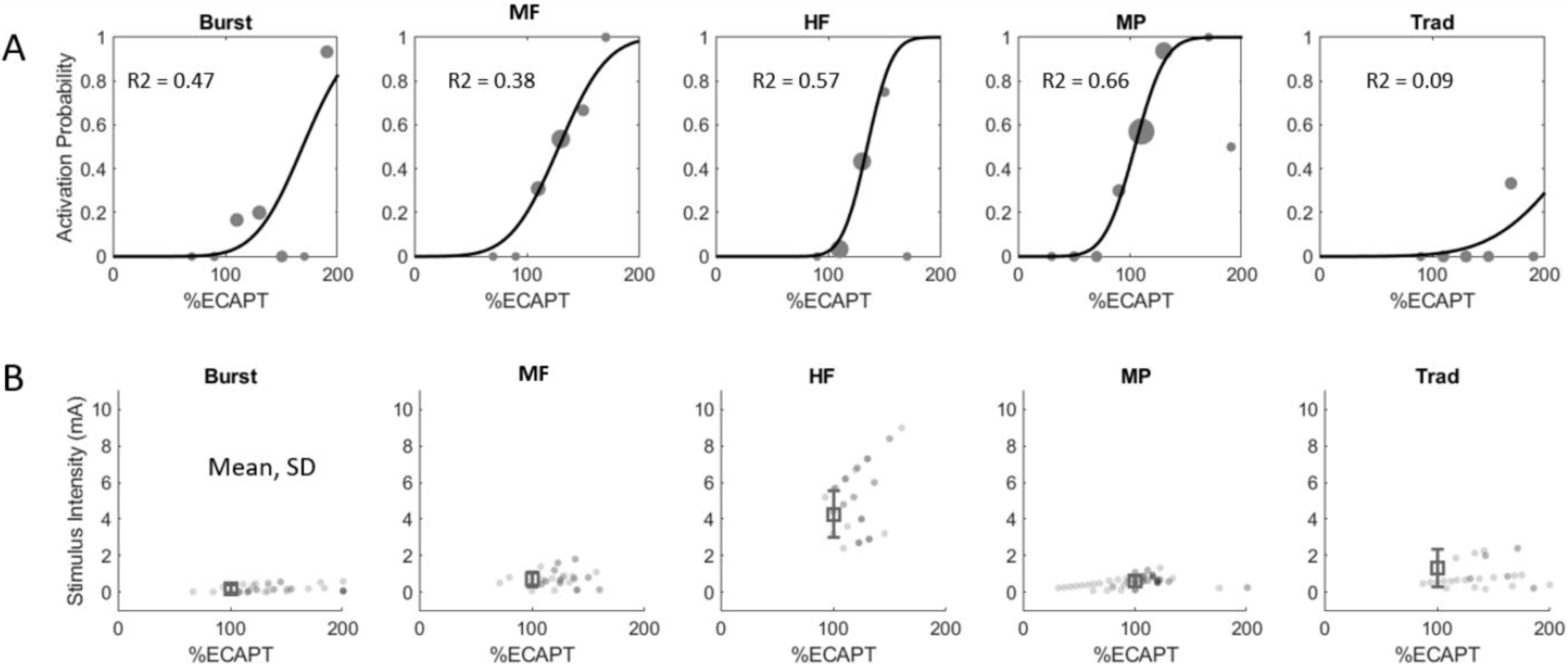
A, Hemodynamic response functions (DSVB activation probability versus SCS amplitude normalized to eCAPT) for the different SCS therapy modes derived from the raw data using a linear mixed-effects model. B, Raw eCAP measurements for each therapy mode consolidated across animal subjects.

### 3.4 Hyperemia magnitude does not differ across clinical therapy modes

The activation functions fit to the DSBV measurements (**Fig. 5A**) were used to estimate the hyperemia magnitude as a function of SCS amplitude for each therapy mode (**Fig. 6A**). All therapies apart from traditional had a positive slope indicating an increase in hyperemia magnitude with increased SCS amplitude. This result is consistent with the eCAP recordings collected during the fUS scans which typically increased in magnitude at higher amplitudes, indicative of greater recruitment of dorsal column fibers (data not shown). Although eCAP amplitude also increased with SCS amplitude for traditional stimulation, only a few significant hemodynamic responses were detected across all experiments. As a result, the estimation of hyperemia magnitude for traditional therapy was not significantly different from control (p = 0.84). The hyperemia magnitudes were quantified by fitting a LME model with therapy mode and SCS amplitude taken as fixed effects (see Methods for details). The expected hyperemia magnitudes estimated by the LME model at 130% eCAPT are shown for each therapy mode in the bar plot in **Figure 6B**. Apart from traditional , all therapy modes at 130% eCAPT drove hyperemia responses significantly greater than control (no SCS) with an average DSBV in the range of 15-20% relative to baseline (note: MF was marginally significant with P < 0.75). When comparing across therapy modes no significant difference was found for hyperemia magnitude. (Note that although traditional therapy clearly showed reduced magnitudes in the few fUS measurements with significant hyperemia, it was not significantly different from other therapy modes in this analysis [or control – no SCS] due to the poor quality of the model fit.) The finding that hyperemia magnitudes were not different between therapy modes is important because it suggests that the clinically-motivated method of scaling SCS amplitude by eCAPT used in this study is appropriate for between therapy comparisons. Specifically, all therapy modes (excluding traditional) drove fUS responses of a similar magnitude allowing for a straight-forward comparison between activation patterns.

**Figure 6.**
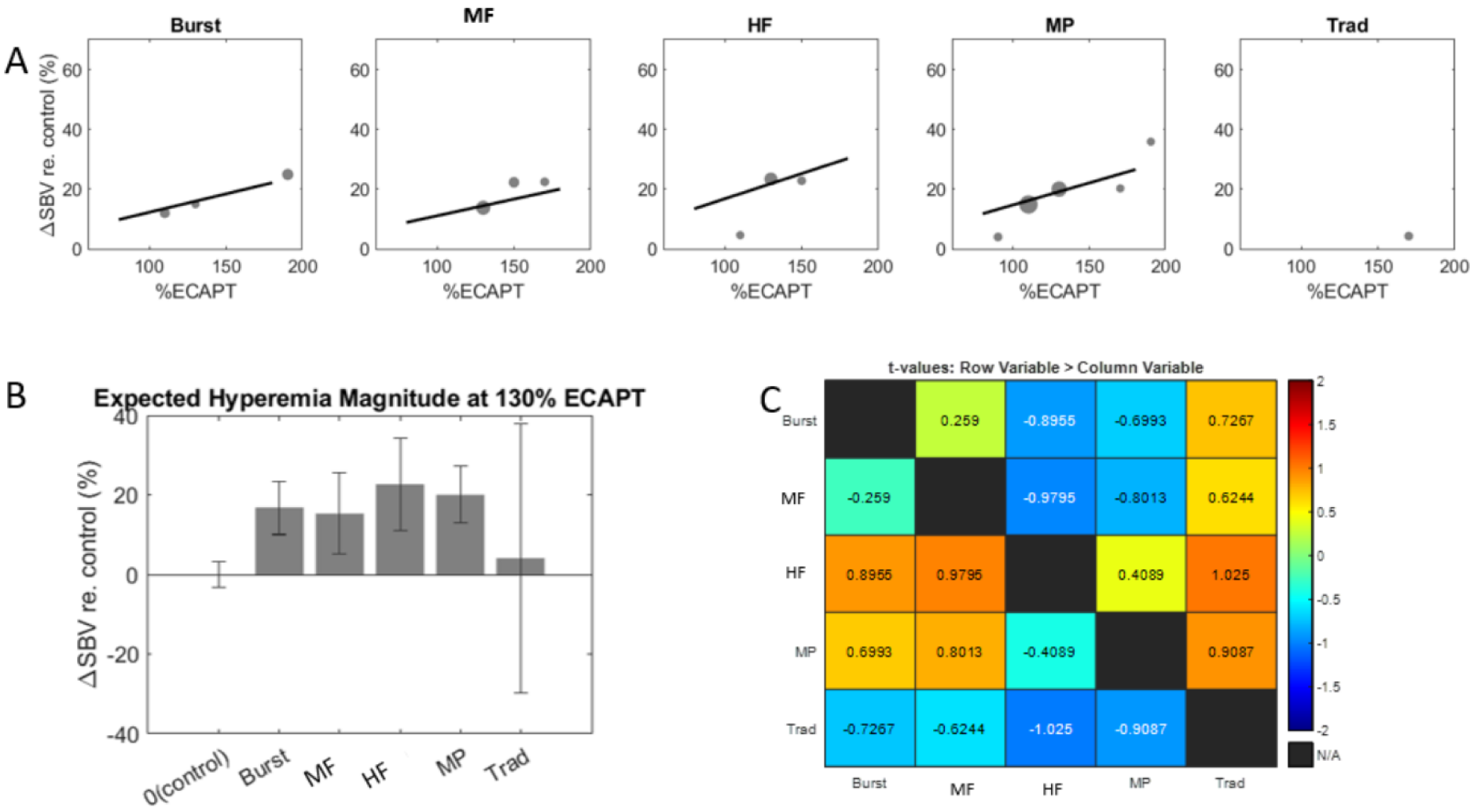
A, Hyperemia magnitude versus SCS amplitude for each therapy mode calculated with the activation functions shown in Figure 5. B, Hyperemia magnitudes for each therapy mode at 130% eCAPT obtained using the linear mixed-effects model. C, t-tables showing all pair-wise comparisons of hyperemia magnitudes. Note that no pair-wise comparisons were statistically significant, all |t| < 2.

### 3.4 Area of hyperemia varies across clinical therapy modes

The next feature considered was the %Area of hyperemia, defined as the percentage of SC pixels (voxels) with significant hyperemia relative to the total number of pixels in the scan area containing the SC. This feature is expected to be highly correlated with neuromodulation area or coverage produced by SCS based on previous work demonstrating afferent somatotopy in the SDH [5]. **Figure 7A** shows model estimates of the %Area of hyperemia as a function of SCS amplitude for each therapy mode. Similar to the results for magnitude, the estimate for traditional had a low correlation with the data (due to a small number of significant responses) while the other therapies were well-fit using a linear function with positive slope. In contrast to magnitude, %Area estimates for both MP and MF had steeper slopes than other therapy modes indicating a more rapid rise in activation coverage with SCS amplitude. The %Area of hyperemia were similarly quantified by fitting a LME model with therapy mode and SCS amplitude taken as fixed effects. At 130% eCAPT the model predicted greatest %Area for MP followed by MF, Burst, HF and Traditional (**Fig. 7B**) with only MP and MF having a %Area (∼10% average coverage) significantly greater than control. The rightmost panel shows the t-values from pair-wise comparisons. Although MP and MF were not significantly different from each other, only MP had significantly greater %Area than HF and Burst. None of the other pair-wise comparisons were statistically significant.

**Figure 7.**
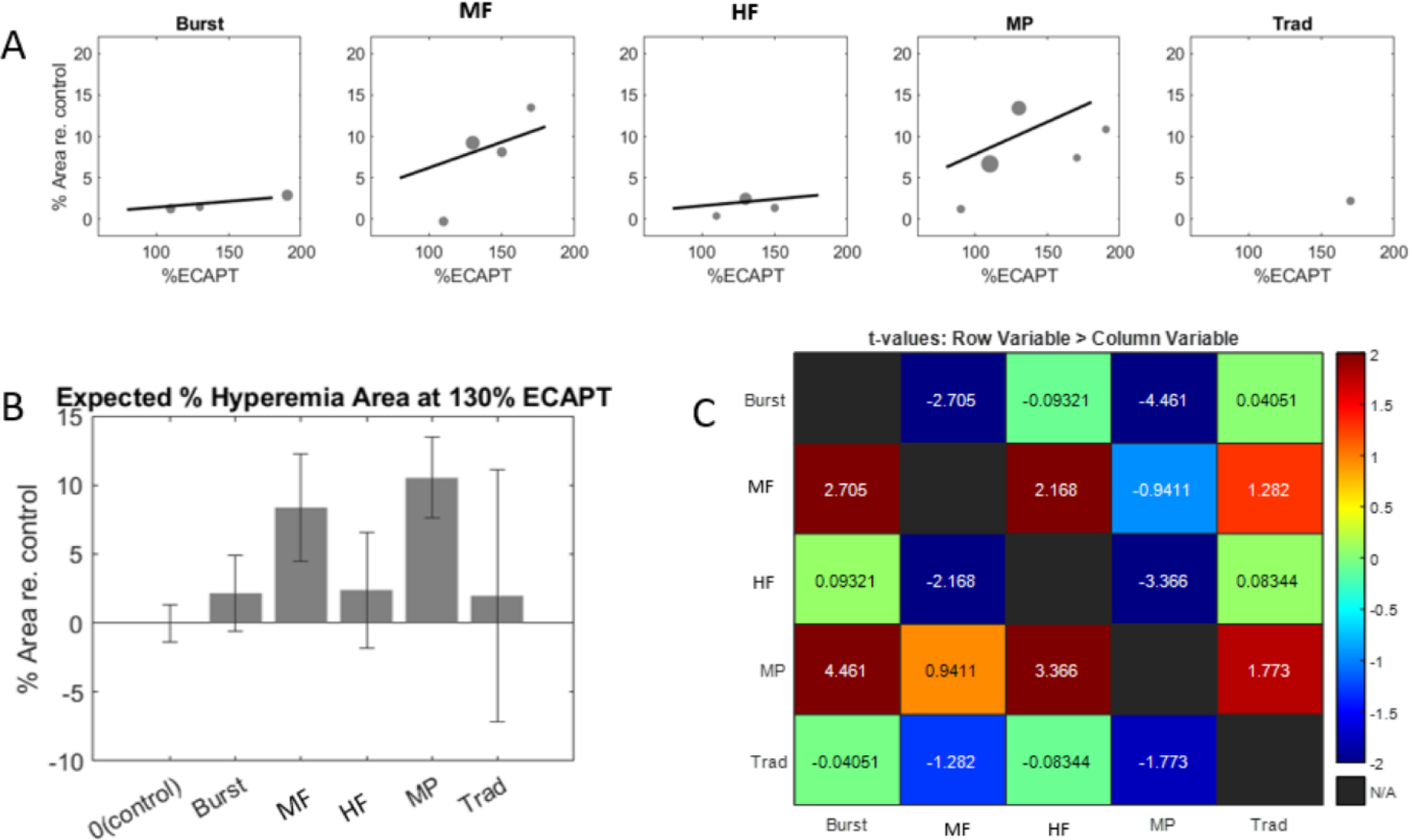
Hyperemia area versus SCS amplitude for each therapy mode calculated with the activation functions shown in Figure 5. B, Hyperemia areas for each therapy mode at 130% eCAPT obtained using the linear mixed-effects model. C, t-tables showing all pair-wise comparisons of hyperemia areas. Statistically significant pair-wise comparisons were determined by |t| > 2.

### 3.5 Depth of hyperemia varies across clinical therapy modes

The final activation feature considered was the Depth of hyperemia, defined as the 75^th^ percentile of the distribution of DSBV depths relative to the dorsal surface of the SC. This feature is expected to be highly correlated with neuromodulation depth or the ability of SCS therapy to target deep neural structures in the SC [32]. **Figure 8A** shows model estimates of the Depth of hyperemia as a function of SCS amplitude for each therapy mode. All therapy modes (excluding traditional) were well-fit using a linear function with positive slope. MP stood out with the steepest slope, indicating activation at greater depths in the SC compared to other therapies over a comparable range of SCS amplitudes. A LME model was again used to quantify depth of hyperemia with therapy mode and SCS amplitude taken as fixed effects. Apart from traditional, all therapy modes activated the SC at depths significantly greater than control. The model estimates are again shown at 130% eCAPT to aid comparison across therapy modes (**Fig. 8B**). The model predicted greatest activation depth for MP (∼1.8 mm below the dorsal surface) followed by MF, Burst, HF and Traditional. Pair-wise comparisons revealed that MP activated the SC at significantly greater depths than all other therapy modes. Although a trend was noted suggesting HF activated the SC at shallower depths relative to Burst and MF, neither this nor any other differences were statistically significant.

**Figure 8.**
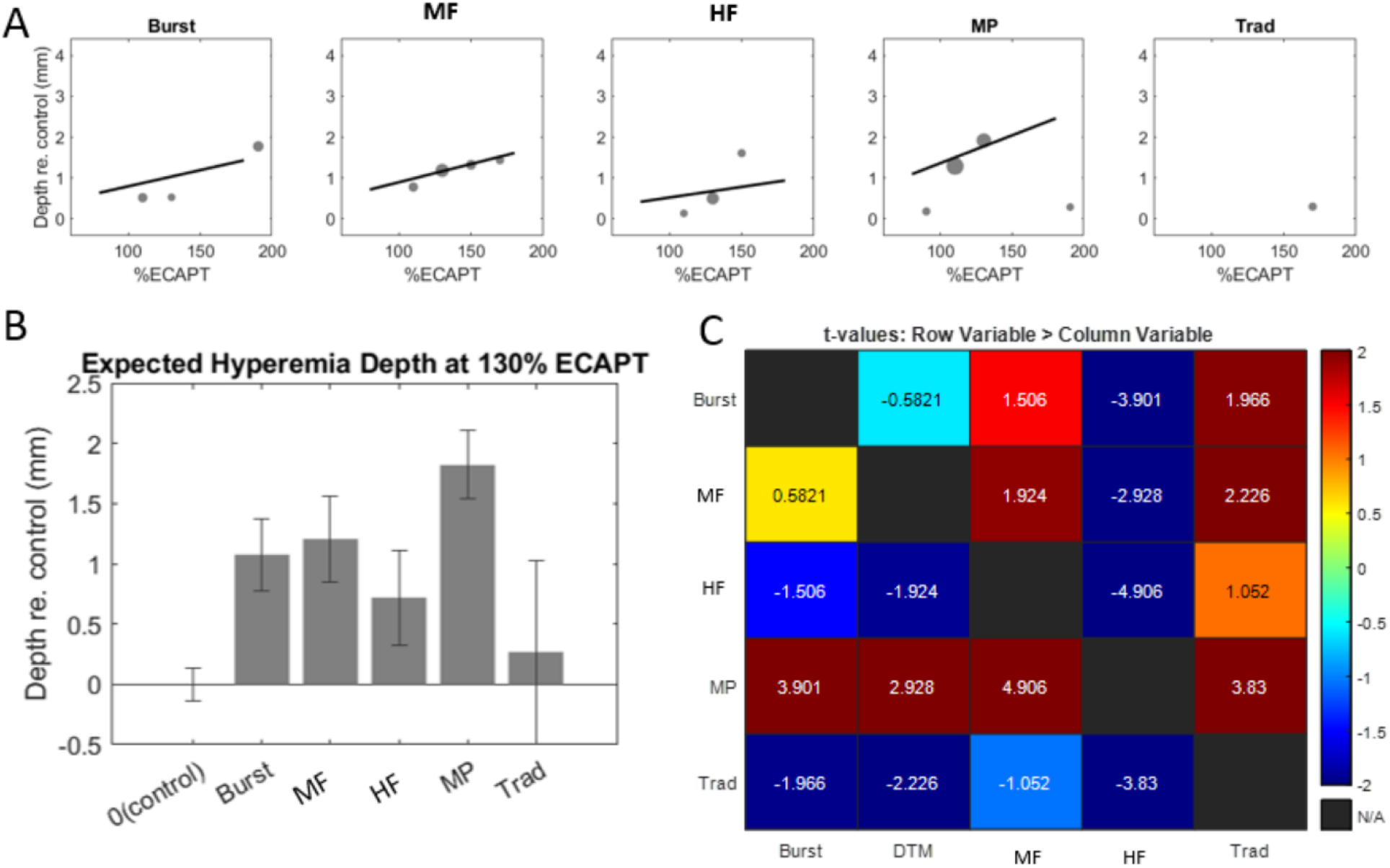
Hyperemia depth versus SCS amplitude for each therapy mode calculated with the activation functions shown in Figure 5. B, Hyperemia depths for each therapy mode at 130% eCAPT obtained using the linear mixed-effects model. C, t-tables showing all pair-wise comparisons of hyperemia depths. Statistically significant pair-wise comparisons were determined by |t| > 2.

## DISCUSSION

### 4.1 fUS reveals neural activation of the SC during SCS

The overlying objective of this study was to measure activation of the spinal cord during clinical SCS in a large animal model that enabled use of clinical hardware. A secondary objective was to compare SC responses across different therapy modalities presently approved to treat neuropathic chronic pain. We developed an acute ovine model using IV anesthesia, low isoflurane and local epidural block that allowed implantation of a standard octal SCS lead. We detected significant hemodynamic responses at levels slightly above dorsal column activation threshold (eCAPT). This novel preclinical model provides new information about neural activation due to the strong neurovascular coupling in the SC [1–3, 5, 33]. Furthermore, it provides a new tool to directly assess neuromodulation during clinically similar SCS and allows comparison of ‘neuromodulation patterns’ within the dorsal horn between clinical therapy modes which was previously not possible.

### 4.2 Hemodynamic responses vary across clinical SCS therapies

Function ultrasound was used to measure differences in neural activation patterns across clinical SCS therapies. We compared traditional low frequency, burst, multifrequency, multiphase and high frequency therapies. The absolute magnitude of activation was clearly weaker for traditional SCS, which only evoked significant responses in a few subjects and required high amplitudes (>150% eCAPT). Both fUS threshold (re eCAPT) and response magnitude elicited by other therapy types were significantly greater than control though not significantly different from each other. Magnitude was evaluated from pixels that demonstrated significant stimulus evoked responses and significant differences were evaluated across the range of tested levels. fUS measures hemodynamic changes in DSBV which in general is not a linear function of SCS amplitude (or of underlying neural spike rate) [3]. Vascular structure and hemodynamics set a limit on the range of responses, which ultimately saturate at high amplitudes [37]. Therefore, there is a theoretical limit to the range over which DSBV faithfully reflects changes in the spike rate of the underlying neural population. This likely impacts the ability to detect differences in SC responses at high amplitudes. In the present study we observed that response magnitude consistently increased with increasing stimulus amplitude. This suggests that response saturation did not have a dominant impact on the results. In summary, the results demonstrate that significant responses can be measured from a broad set of SCS therapies and support the use of eCAPT to normalize across-therapy comparisons.

In contrast to response magnitude, the area of activated regions in the SC varied significantly across therapy modes. Activation area was computed by adding the number of significantly activated pixels, regardless of activation magnitude. MP and MF activated a significantly larger SC area relative to control. Pairwise therapy comparisons revealed MP activated a greater area relative to HF and burst. All other pairwise comparisons were not significant. The statistical results reflect the finding that activation areas for some therapies were highly variable across subjects. For example, MF activated distinct areas in some subjects (e.g. **Fig. 4E**) while driving no significant responses in others (e.g. Subj#107-070, data not shown). Although the average hyperemia area was larger for MP than MF, the high variability of MF made statistical comparisons difficult. Due to strong neurovascular coupling in the SC, it is reasonable to assume a larger area of hemodynamic response results from a greater volume of neural activation. This suggests that MP therapy activates a greater volume of neural tissue than the other therapies examined.

All therapies (apart from traditional) evoked significant hyperemia at depths ranging on average from ∼0.5-2 mm below the dorsal surface (at SCS amplitudes of 130% eCAPT). Hyperemia depth had the most prominent differences across therapy type. MP activation reached to significantly greater depths compared to any other therapy tested. Burst and MF activated the next greatest depths on average although neither differences between these therapies nor between traditional and HF were statistically significant. The neurovascular coupling assumptions noted above suggest that MP therapy modulates neurons deeper in the SC than other therapy modes. The average hyperemia depth of MP was ∼1.8 mm which would cover approximately dorsal horn laminae I-V in the sheep and thus encompasses all pain processing laminae I-III (spinal level T13-L1, [38]). Assuming a similar penetration depth, MP would also cover the pain processing laminae in the human dorsal horn [39].

### 4.3 Comparison with previous studies

The finding that MP therapy provides a greater area of coverage and depth of response versus other common stimulation types is consistent with previous studies and the developmental objectives of this therapy. The BENEFIT-01 clinical study investigated the impact of a wide range of SCS parameters on patient perception [40]. Key findings of this study include that perception threshold (PT) is primarily influenced by pulse width, but not pulse frequency, and that 14 mm spacing between active electrode centers results in lower thresholds than closer spacing (eg 7 mm). Other studies have demonstrated that PT closely corresponds to threshold for neural activation of dorsal column fibers [11, 41], thereby implying that parameter changes that decrease PT also decrease neural activation threshold and by extension decrease threshold for effective subthreshold neuromodulation [32].

The findings of previous studies were incorporated into computational models and combined with known spatiotemporal integration properties of neural circuits to optimize MP therapy parameters. The therapy development objective was to achieve efficient neuromodulation of a broad area of the SC targeting deep gray matter regions (ie dorsal horn). Computational modeling of the optimized MP therapy demonstrated that the stimulation pattern achieved these objectives more efficiently (using less power) compared with other common stimulation patterns [42]. The findings of the current study are consistent with these previous model predictions and serve to further validate the MP approach.

A recent fUS study in the rat SC demonstrated high resolution spatiotemporal responses of spinal nociceptive circuits in both normal and inflammatory animal models [5]. Robust hemodynamic responses to stimulation were observed with a time course consistent with the current study. In contrast, the rat study primary found hyperemia (DSBV increases) in response to stimulation while the present study found that both hyperemia and congestion (DSBV decreases) were common. There are many methodological differences between the studies that may account for this discrepancy. First, the present study used direct SCS while the previous used peripheral nerve stimulation. Second, the present study used a large animal model with a clinical grade SCS system while the previous study was conducted in the rat using custom stimulation hardware. Finally, although both studies utilized low concentrations of isofluorane, which is known to diminish neurovascular coupling [43], there were several differences in I.V. anesthetics which may have impacted the responses.

Other studies used fUS to measure SC hemodynamic responses to SCS using both large (pig) and small (rat) animal models [4, 44]. Consistent with both the present study and the Claron study, the authors found both hyperemia and congestion were common in pigs but hyperemia was most common in rat. At present it unclear how to interpret congestion and why it might be more prevalent in large animal models. We found that congestion and hyperemia magnitudes (not area and depth) are correlated and in addition that voxels do not switch from positive to negative DSBV as SCS parameters are changed. These results suggest an underlying anatomical substrate which reflects changes in neural activity. One hypothesis is that congestion reports net inhibition (a decrease in FR of the neural population). Another is that decreases in blood volume are due to vasoconstriction of larger arteries and veins by direct SCS activation of smooth muscle cells. If true, differences in congestion in large and small animals could be due to differences in smooth muscle thresholds relative to dorsal column fiber threshold. Future experiments using methods to selectively inhibit interneurons (i.e. piezoelectric cooling) or pharmacologically induce vasoconstriction may resolve these possibilities.

### 4.4 Implications for SCS mechanisms of action

Functional ultrasound imaging demonstrated significant spinal blood volume changes in response to SCS. Changes were seen underlying the active cathodes from the most dorsal aspect near the PSV (dorsal columns) to a depth of several mm in the ventral direction (gray matter of the DH). This suggests the following mechanisms of action: 1. Suprathreshold SCS induces DSBV in the white matter due to extensive vascular innervation there (e.g. anterospinal artery and posterior spinal vein) which circulates blood to the underlying gray matter. The main blood supply for the dorsal columns likely arises near their cell bodies in the dorsal root ganglion. 2. SCS can induce activation of the dorsal horn neurons at similar amplitudes, inducing an increased DSBV due to neurovascular coupling causing an increased blood supply. 3. SCS may also induce indirect activation of the DH via antidromic activation of the DCFs. The response of the DH may be excitatory or inhibitory depending on the balance of excitation and inhibition in the specific region under investigation. 4. As DSBV was only seen for suprathreshold SCS in this study, the mechanisms of action for subthreshold SCS may be different than those reported here. However, suprathreshold changes in blood flow may still serve to highlight the neuronal locations that would be most engaged by subthreshold stimulation. Future studies using synaptic blockers may help to elucidate contributions of direct versus indirect activation of the spinal gray matter.

### 4.5 Study Limitations

Although the present study has provided new insights to SC activation during clinical SCS, there are several limitations that could be addressed in future studies. The study was conducted in naive animals under anesthesia and analgesic agents. Although isofluorane anesthesia was limited to 1% or below, this may have impacted the hemodynamic responses (but not the therapy comparisons, due to randomization of the test sequences and mixed-effects analysis). In addition, analgesic agents including fentanyl were administered for pain control as necessary for the invasive laminectomy procedure but this might have impacted nociceptive responses as measured by fUS through neurovascular coupling. As a result of these experimental manipulations, the spontaneous activity of both the DC fibers and gray matter was probably very low which may have been why suprathreshold SCS was required to detect significant hemodynamic responses.

Future studies using neuropathic pain models [45] and/or chronic recording methods [38] [46] could be developed to study animal models more closely representative of human neuropathic pain. Such models should theoretically have hyperactivity in neuropathic sites on the SC. The present fUS technology may resolve such sites by comparing baseline signal strength to healthy control regions. The neuropathic sites could be monitored during subthreshold SCS to detect changes in hyperactivity and its time course compared with human clinical findings, where maximum pain relief from subperception SCS typically takes hours to days. This model would be a powerful tool to investigate parameters that influence pain relief and help to guide improvements in SCS therapy efficacy.

## CONCLUSIONS

This work demonstrates that fUS can effectively measure SCS neural response patterns in the pain processing laminae of a large animal model implanted with a clinical SCS system. Hemodynamic responses in the SC varied significantly across SCS therapy modes, with Multiphase stimulation providing a greater area of coverage and depth of response versus other common stimulation types.

## Authorship Statement

Koeun Lim, Sean Slee and Andrew Kibler were involved in the study design, data collection, and data analysis. All authors had access to relevant data, were involved in interpreting the findings, and participated in drafting, reviewing, and approving this manuscript.

## ETHICS STATEMENT

Experimental procedures were approved by the Legacy Research Institute Animal Care and Use Committee. The animals we cared for according to the USDA Animal Welfare Act standards and the eighth edition of *The Guide for Care and Use of Laboratory Animals*.

## ACKNOWLEDGEMENTS

The authors gratefully acknowledge expert technical assistance from Dan Dahlke and Courtney Fuller and thank the staff at Legacy Research Institute for exceptional animal care.

## ABBREVIATIONS

SCS: Spinal cord stimulation
fUS: Functional ultrasound
ΔSBV: changes in flowing spinal blood volume
eCAP: evoked compound action potential
EMG: electromyographic
SDH: superficial dorsal horn
eCAPT: eCAP threshold
DC: dorsal column
IV: intravenous
CRI: constant rate infusion
bpm: breaths per minute
EPG: external pulse generator
PSV: posterior spinal vein
ms: milliseconds
BST: Burst
MF: Multi-frequency
HF: High kHz frequency
MP: Multiphase
Trad: Traditional
ROI: region of interest
ROIh: hyperemia region of interest
ROIc: congestion region of interest
SD: mean (*μ̑*), variance (*σ̑*^2^), standard deviation
LME: linear mixed-effects model

